# Early adversity changes the economic conditions of structural brain network organisation

**DOI:** 10.1101/2022.06.08.495303

**Authors:** Sofia Carozza, Joni Holmes, Petra E. Vértes, Ed Bullmore, Tanzil M. Arefin, Alexa Pugliese, Jiangyang Zhang, Arie Kaffman, Danyal Akarca, Duncan E. Astle

## Abstract

Early adversity can change educational, cognitive, and mental health outcomes. However, the neural processes through which early adversity exerts these effects remain largely unknown. We used generative network modelling of the mouse connectome to test whether unpredictable postnatal stress shifts the constraints that govern the formation of the structural connectome. A model that trades off the wiring cost of long-distance connections with topological homophily (i.e. links between regions with shared neighbours) generated simulations that replicate the organisation of the rodent connectome. The imposition of early life adversity significantly shifted the best-performing parameter combinations toward zero, heightening the stochastic nature of the generative process. Put simply, unpredictable postnatal stress changes the economic constraints that shape network formation, introducing greater randomness into the structural development of the brain. While this change may constrain the development of cognitive abilities, it could also reflect an adaptive mechanism. In other words, neural development could harness heightened stochasticity to make networks more robust to perturbation, thereby facilitating effective responses to future threats and challenges.

**Significance statement:** Children who experience adversity early in life – such as chronic poverty or abuse – show numerous neural differences that are linked to poorer cognition and mental health later in life. To effectively mitigate the burden of adversity, it is critical to identify how these differences arise. In this paper, we use computational modelling to test whether growing up in an impoverished and unpredictable environment changes the development of structural connections in the mouse brain. We found that early adversity appears to introduce more stochasticity in the formation of neural architecture. Our findings point to a potential mechanism for how early adversity could change the course of child development.

## Introduction

The structure of the human brain undergoes complex changes over the first three decades of life^1^. At the macroscopic level, neural development proceeds through the formation of a network of white-matter projections between populations of neurons, a process both subject to genetic control and environmental regulation^2–4^. A complete wiring map of the brain, known as a “connectome”, can be reconstructed through diffusion-weighted magnetic resonance imaging (MRI) and analysed using graph theory^5^. Healthy neural architecture is characterised by a precise pattern of organisation, or topology, that emerges over the course of childhood^6,7^. For instance, brain networks exhibit small-worldness, a balance between a short average path length and high clustering that permits both integrated and segregated processing of information^8,9^. Features of connectome organisation can predict developmental differences across individuals, including variation in cognitive ability and mental health^10,11^.

The structural organisation of the brain emerges amid a tight set of constraints. The most significant of these is the biophysical embedding of the network, because of which long-distance connections incur a large metabolic cost^12^. The brain has adapted to limit this cost by making parsimonious use of energy and space, creating comparatively expensive features – such as connections between spatially distant regions – judiciously^9,13^. But cost minimisation alone cannot account for the observed organisation of biological neural networks^14,15^. Rather, the brain appears to negotiate an economic trade-off between the physical cost of structural connections and the topological value they add to the network^9,16,17^. Recent advances in computational modelling offer a way to directly investigate the constraints that govern the development of the connectome by generating networks using different wiring rules^14,16,18–20^. Studies employing this approach have shown that slight manipulations in the trade-off between two key generative model terms - wiring cost and topological value - can reproduce real-world diversity in structural brain organisation, and account for differences in behavioural phenotypes^16,21,22^. However, we do not yet know which developmental factors, including social environmental conditions in early life, modulate the wiring economy and thus shape the trajectory of brain network development.

The quality of the early environment is a critical determinant of neurodevelopment^23^. Children who experience adversity or maltreatment show subtle differences in the organisation of their connectomes, including lower connectivity between modules and altered centrality of regions such as the amygdala^24,25^. Such neural differences may be conducive to navigating a hostile and unpredictable early environment, but may come at the expense of poorer cognition and mental health later in life^26^. Due to the methodological and ethical limits of human research, experimental studies in rodent models have proven invaluable for establishing the causal role of adversity in neural outcomes^27^. Recent work in mice has shown that early-life stress causes local changes in brain network organisation, including an increase in frontolimbic connectivity and decrease in efficiency of the amygdala, that drive a global increase in small-worldness and heightened anxiety-related behaviour^28,29^. The increasingly thorough demonstration of adversity-related differences in brain structure highlights a crucial mechanistic gap in our understanding: how does early adversity alter the development of network-level brain organisation?

In the current study we test whether early adversity alters the wiring economy of the developing mouse connectome using a paradigm of unpredictable postnatal stress (UPS). UPS pups are raised under conditions of limited bedding to mimic impoverishment and are also exposed to unpredictable hour-long bouts of maternal separation and nest disruption to model chaotic and complex adversity^28,30^. We reconstructed the structural connectomes of 49 adult mice, half of which were exposed to UPS during the first four weeks of life^30^. Using generative network modelling, we computationally simulated realistic networks for each animal and evaluated how well they replicated the observed connectomes. We then tested for differences in the economic conditions of brain development by comparing the generative model parameters that most closely replicated the connectomes of each group. Finally, we explored the developmental implications of shifts in the wiring economy of the brain.

## Results

### Empirical connectomes

At birth, N = 49 pups were randomly assigned to a control or UPS^30^ condition (**Figure 1a)**. Mice were kept in rearing conditions until adolescence and sacrificed in adulthood, at which point diffusion imaging was performed (see Methods). Using probabilistic tractography, we reconstructed binary structural connectomes for each mouse. The connectomes showed no differences between groups on gross measures of global topology, including on number of edges (*p* = 0.89), number of long-distance connections (*p* = 0.52), maximum modularity (*p* = 0.72), global efficiency (*p* = 0.71), or small-worldness (*p* = 0.47) (see **Methods**; **Supplementary Table S1**). Groups did not differ on the distributions of key local characteristics, including node degree, clustering coefficient, betweenness centrality, edge length, mean matching index, and nodal efficiency (all *p* > 0.96) (see **Methods**; **Supplementary Table S2**).

**Figure 1.**
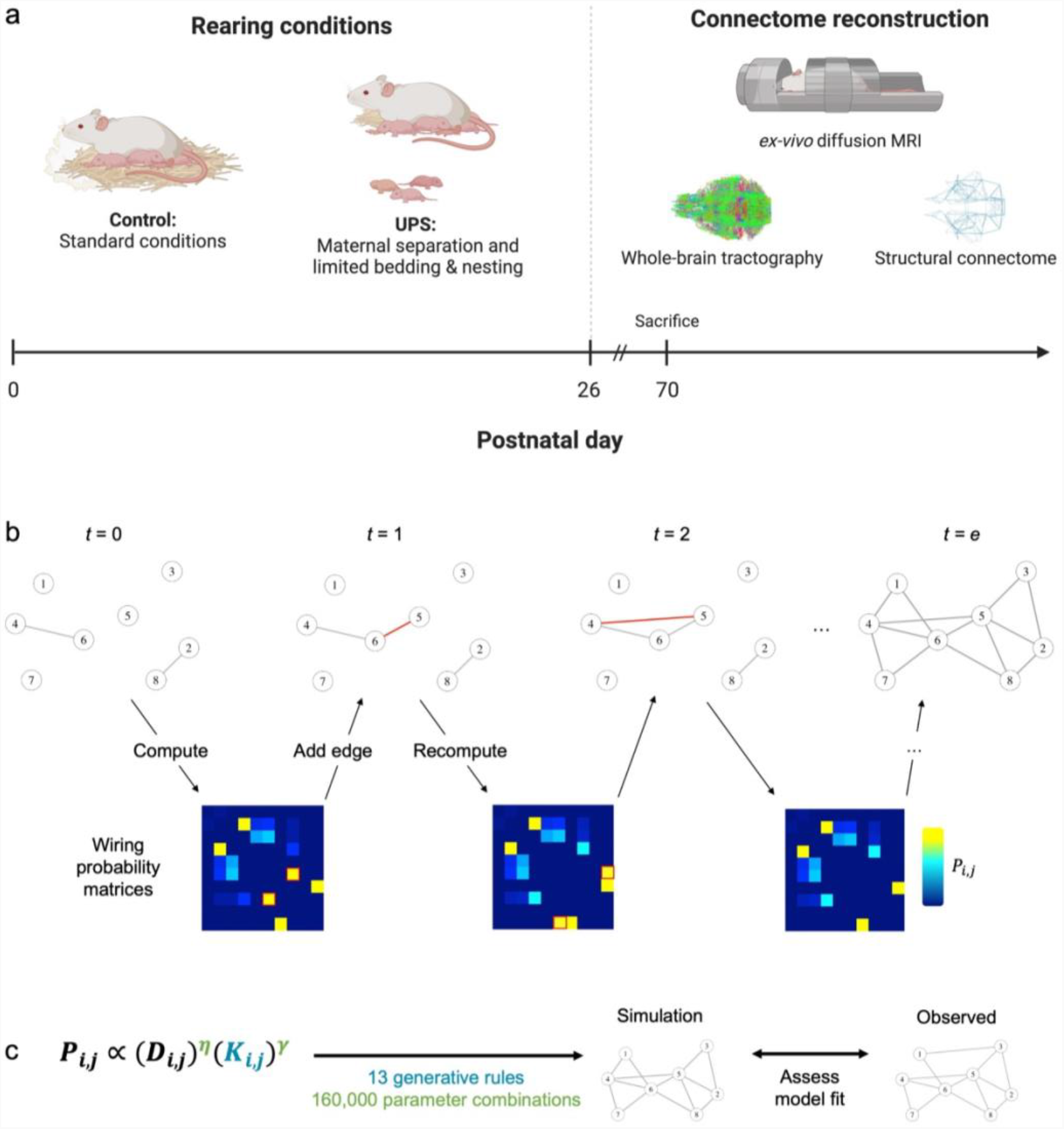
Experimental design and generative modelling procedure. **(a)** On postnatal day 0, N = 49 pups were randomly assigned to a paradigm of unpredictable postnatal stress or standard rearing conditions until postnatal day 26. After postnatal day 70, mice were sacrificed and ex-vivo diffusion imaging was performed. Whole-brain probabilistic tractography was used to reconstruct the structural connectome of each animal. **(b)** An illustration of the generative process using a simplified connectome of ten nodes. Starting from a sparse seed network (t = 0), edges are added one at a time until the simulation reaches the number of edges found in the observed connectome (t = e). The matrix of wiring probabilities is updated at each step, allowing for dynamic shifts as the topology of the network emerges. **(c)** By systematically varying generative rules and parameter combinations, it is possible to identify the topological term K and the parameters *η* and *γ* that best simulate the organisation of the observed connectome.

### Generative modelling procedure

To simulate the formation of each connectome, we formalised a trade-off between two competing factors: the wiring cost incurred by new connections and the topological value they add to the network^16,21^. The cost term penalises long-distance connections, thereby capturing the evolutionarily conserved drive to minimise the metabolic and material expense of axonal projections^9,18^. The value term favours connections between regions that share some topological property, such as a similar pattern of clustering or a large number of existing connections^7,9,18^.

The model simulates connectome formation by incrementally adding connections, one at a time, from some initial conditions. At each step, it estimates the likelihood of potential new structural connections using a simple probability equation^16,21^:

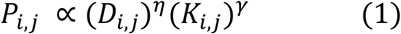

where *P*_*i,j*_ is the probability of forming a binary connection between any two previously unconnected regions of the brain, *i* and *j*. The first term *D*_*i,j*_ represents the wiring cost. As the resources required by an axonal projection increase with its length^9^, *D*_*i,j*_ approximates the cost of a connection using the Euclidean distance between the brain regions it would connect. The term is scaled by a parameter *η*, which determines the strength of its contribution to the overall wiring probability. To penalise longer distance connections, *η* is negative.

The second term *K*_*i,j*_ represents the topological value of a connection and can take numerous forms. Following previous work^16,21,22^, we tested thirteen variations of *K* (known as “generative rules”), each one quantifying a different topological relationship between the two nodes *i* and *j*. The generative rules we considered fall in three categories: (i) homophily models, which favour connections between nodes with similar connectivity neighbourhoods; (ii) clustering-based models, which utilise the clustering coefficients of the regions; and (ii) degree-based models, which utilise their node degree. *K*_*i,j*_ is scaled by a parameter *γ*, which is positive to favour connections with a higher topological value.

At every step of the generative process, the model multiplies the cost and value terms for each pair of regions to produce a matrix of relative wiring probabilities, and probabilistically chooses a “winning” edge to add to the simulation (**Figure 1b**). Given that every new connection changes the topology of the network, and therefore also the *K*_*i,j*_ term for some possible connections, the model iteratively updates *P*_*i,j*_ over time. In other words, the model continually re-computes the probability of future connections.

Shifts in wiring probability can occur quite rapidly, especially whilst the connectome is sparse^22^. For example, consider the network at step *t* = 0 in **Figure 1b**. Suppose it is growing according to a generative rule that favours connections between regions with shared neighbours. According to the probability function (Eq. 1), nodes 4 and 5 would be unlikely to wire together at first, because they are relatively distant and share no neighbours. Instead, at step *t* = 1, a connection forms between proximal nodes 5 and 6. However, this new connection gives nodes 4 and 5 a shared neighbour and therefore increases the topological value of forming a direct connection, which occurs at step *t* = 2, despite the greater distance between them. Whilst wiring cost remains the same across development, the topological value of connections, and therefore the overall wiring probability, is dynamic from one step to the next. As the network grows, longer connections become increasingly likely as the topological value added by new links outweighs the penalisation of wiring cost^31^.

The generative process terminates when the synthetic network reaches the number of edges of the connectome that the model is simulating. By varying the generative rule used as the topological term *K*_*i,j*_, and the *η* and *γ* parameters, it is possible to systematically manipulate the conditions that govern the development of the synthetic network (**Figure 1c**). Identifying the rules and parameters that best simulate the real connectomes of individuals can thus shed light on what may be guiding their structural neurodevelopment^18,21,22^.

### Homophily-based simulations achieve best model fit

We first sought to identify the generative rule that most successfully reproduced the structural connectomes of our sample of mice (N = 49). For each animal and generative rule, we tested 160,000 parameter combinations evenly distributed throughout the space defined by −10 ≤ *η* ≤ 0 and 0 ≤ *γ* ≤ 10. Beginning with a sparse seed network of edges shared across all animals (see **Methods**; **Supplementary Figure S1**), connections were added according to the probability function (Eq. 1) until the synthetic network reached the number of edges of the empirical connectome of that animal.

At the end of the generative process, we assessed how well each synthetic network fit the connectome it was simulating using the following energy equation^21^:

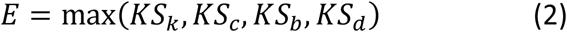

where the terms are the Kolmogorov-Smirnov (KS) statistics comparing the synthetic and empirical distributions of node degree (*k*), clustering coefficient (*c*), betweenness centrality (*b*), and edge length (*d*). These four measures are critical properties of networks that are linked to stress exposure and psychiatric conditions^32,33^ and have previously been used to assess the similarity of empirical and economically simulated connectomes ^21,22,34^. As the energy is the maximum of the four statistics, a lower energy corresponds to better model fit. In other words, Eq. 2 compares the organisation of each synthetic network to the organisation of the biological connectome; if the networks are similarly organised, then the energy will be low.

To assess the performance of the models, we compared the lowest-energy simulation produced by each rule. All generative rules outperformed a purely spatial model that considered only wiring cost (**Figure 2a**; **Supplementary Table S3**). An ANOVA and post-hoc Tukey test confirmed that models specifying homophily as the topological *K*_*i,j*_ term achieved lower energy than those utilising clustering (*diff* = -0.090, *p* = 1.97 × 10^−12^) or degree (*diff* = -0.020, *p* = 1.16 × 10^−9^). Thus, generative models that trade-off the wiring cost of a connection with a measure of neighbourhood similarity produce synthetic networks whose global topological distributions closely resemble those of the observed connectomes. As multiple models achieved low energy, the success of the top-performing models from each category was examined further.

**Figure 2.**
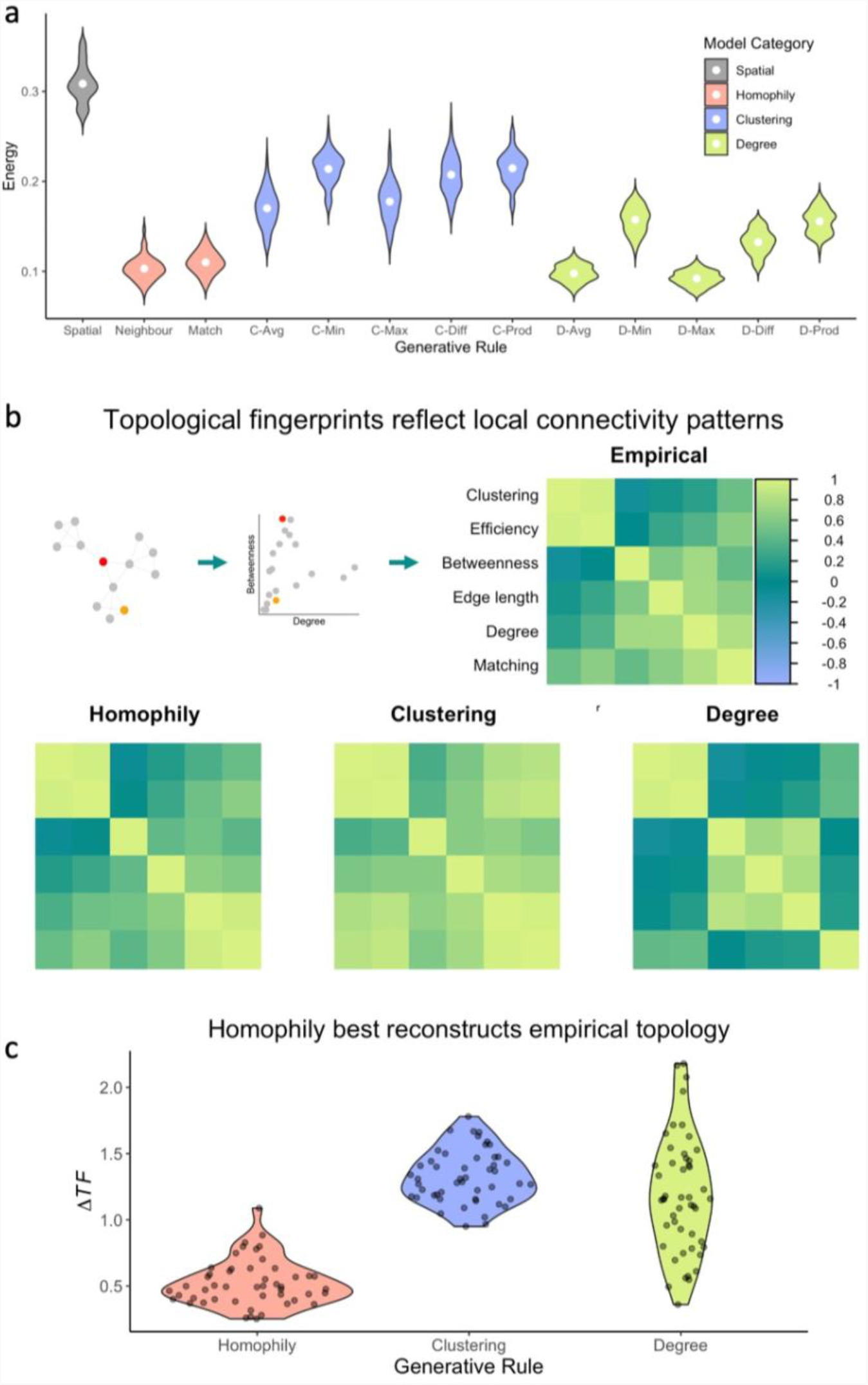
Relative performance of generative network models in replicating the organisation of empirical connectomes. **(a)** The energy of the top-performing synthetic networks for each animal (N = 49) across thirteen generative rules: a purely spatial model, which considers only the distance between two regions; two homophily models, which also consider a measure of the similarity of the neighbourhoods of the respective regions; five clustering-based models, which compare the clustering coefficients of the regions; and five degree-based models, which compare their node degree. White points indicate the sample mean. **(b)** The topological fingerprint is a correlation matrix of local network statistics, including node degree, clustering coefficient, betweenness centrality, total edge length, local efficiency, and mean matching index. Topological fingerprints are shown for the empirical networks and the best-performing rules across the three categories of generative models. Across all four matrices, the value of the correlation can be inferred from the colour bar, which spans -1 (purple) to 1 (green). Correlations shown are the sample average (N = 49). **(c)** Across the sample (N = 49), homophily achieves lowest Δ*TF*, a measure of the discrepancy in local patterns of connectivity between the simulations and empirical connectomes.

### Homophily best recapitulates the local properties of observed networks

The energy equation effectively assesses how closely the statistical distributions of nodal characteristics of the synthetic networks resemble those of the empirical connectomes. However, brain networks also exhibit local patterns of relationships between nodal characteristics. For instance, nodes with high betweenness centrality tend to be lower in clustering, given their position between modules^35^.

To address this, we characterised the local organisational properties of the empirical and simulated networks using a method called the “topological fingerprint,” which has recently been developed for this purpose^31^. First, we selected the lowest-energy simulations produced by each generative rule. We then calculated six common measures of nodal topology, including degree, betweenness centrality, clustering coefficient, edge length, local efficiency, and mean matching index. Next, we computed correlations between these measures, calculated the sample average, and summarised the results in a 6-by-6 matrix.

These matrices are called topological fingerprints because they summarise the unique patterns of local organisation found across a network. Topological fingerprints (TF) for the empirical connectomes and the top-performing generative models from each category can be found in **Figure 2b** (all other rules are shown in **Supplementary Figure S2a**). A visual comparison of the topological fingerprints offers a way of estimating how well the generative models replicate the local properties of the connectomes. To formalise this assessment quantitatively, we also calculated the difference in their topological fingerprints, (ΔTF) according to the following equation^31^, which implements the Euclidean norm:

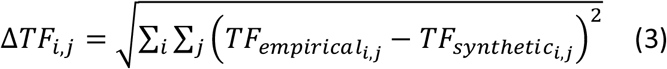

The homophily model achieved the lowest ΔTF, confirming the visual impression that its topological fingerprint was most similar to that of the empirical connectomes (**Figure 2c**; comparable results are shown for all other rules in **Supplementary Figure 2b**). In other words, a model that balances the cost of an additional connection against the number of shared neighbours produces networks with local patterns of organisation that closely resemble those of the rodent connectome, even though local topology was not explicitly optimised by the energy function.

### Homophily replicates spatial layout of empirical networks

Given that the wiring of biological neural networks is shaped by their embedding in anatomical space^36^, realistic synthetic connectomes should ideally exhibit a spatial layout akin to that of connectomes derived from tractography. To test this similarity, we first calculated the six characteristics of each node of the parcellation, averaged across the sample, then correlated the values between simulated and empirical connectomes^21,22^. As shown in **Figure 3**, all four measures included in the energy equation exhibited significant correlations: degree (*r* = 0.360, *p* = 2.68 × 10^−5^), clustering coefficient (*r* = 0.346, *p* = 5.40 × 10^−5^), betweenness centrality (*r* = 0.530, *p* = 9.33 × 10^−11^), and edge length (*r* = 0.543, *p* = 2.52 × 10^−11^). Correlations were also observed between synthetic and empirical nodes on local efficiency (*r* = 0.420, *p* = 7.01 × 10^−7^) and mean matching index (*r* = 0.334, *p* = 1.02 × 10^−4^), confirming that the simulations replicated the spatial layout of nodal features that were not used to optimise model parameterisation.

**Figure 3.**
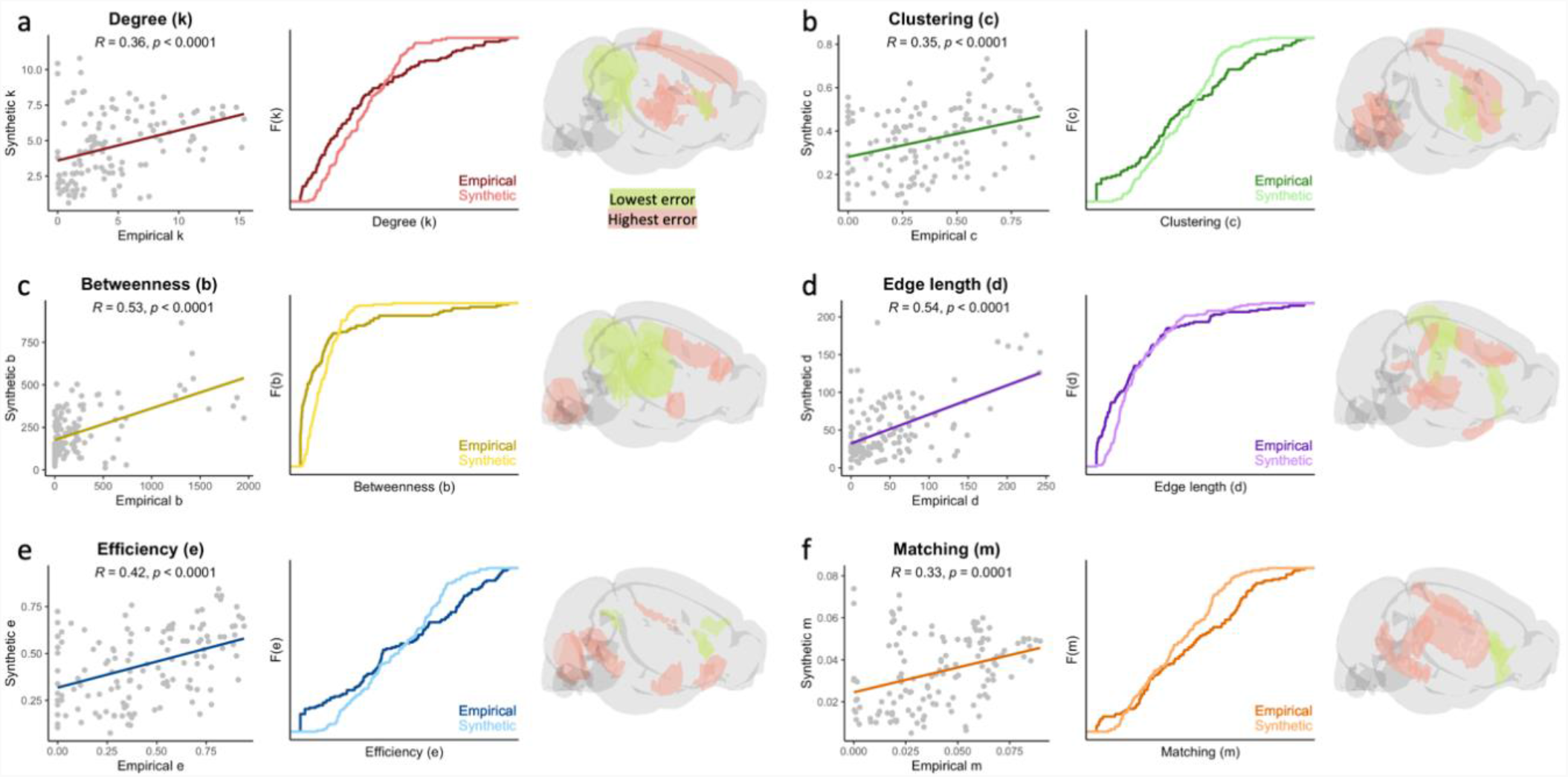
Simulated networks replicate spatial layout of empirical connectomes. Each point in the scatterplots represents the nodal measure for one of the 130 regions of the parcellation, taken as the average value across animals (N = 49). For each of the six measures, a significant positive correlation exists between the nodes of synthetic and empirical networks. A cumulative density function of the measure is also displayed, as well as a visualisation of the mouse brain in which the five regions with the lowest and highest error (i.e., discrepancy between synthetic and empirical networks) are highlighted in green and red respectively. Four of the statistics (**(a)** node degree, **(b)** clustering coefficient, **(c)** betweenness centrality, and **(d)** total edge length) are terms of the energy equation used to assess the fit of the synthetic networks, while the remaining statistics (**(e)** local efficiency and **(f)** mean matching index) are not.

We also assessed discrepancies between the simulated and observed connectomes in the layout of these local characteristics. At each node, we computed a measure of spatial error by subtracting the average value of each characteristic in the synthetic networks from its average value in the empirical connectomes^22^. Thus, a lower spatial error indicates more similarity between the local topology of a particular region in the simulations and in the observed connectomes. While overall spatial error was distributed throughout the brain (**Supplementary Table S4**), a significant correlation was observed between spatial error and node degree in the seed network (*r* = 0.436, *p* = 2.221 × 10^−7^) (**Supplementary Table S5**). This indicates that generative models may benefit from instructions as to where to begin adding connections if they are to best replicate the spatial patterning of network characteristics.

### Early adversity attenuates wiring constraints in optimal simulations

Across all generative models, the homophily model implementing the neighbour rule exhibited the smallest coefficient of variation in the *γ* parameter and second smallest in the *η* parameter (**Supplementary Table S3**). Thus while this rule was best able to account for variations in topology across animals, it did so through minute adjustments in the weighting of its cost and value terms, likely indicative of the highly regulated nature of connectomic organisation (**Figure 4a**). To obtain maximally precise parameters for each animal, we therefore performed a second search of 40,000 parameter combinations in a narrow space centred at the apparent minimum of the energy landscape: −3.75 ≤ *η* ≤ −1.75 and 0.2 ≤ *γ* ≤ 0.6.

**Figure 4.**
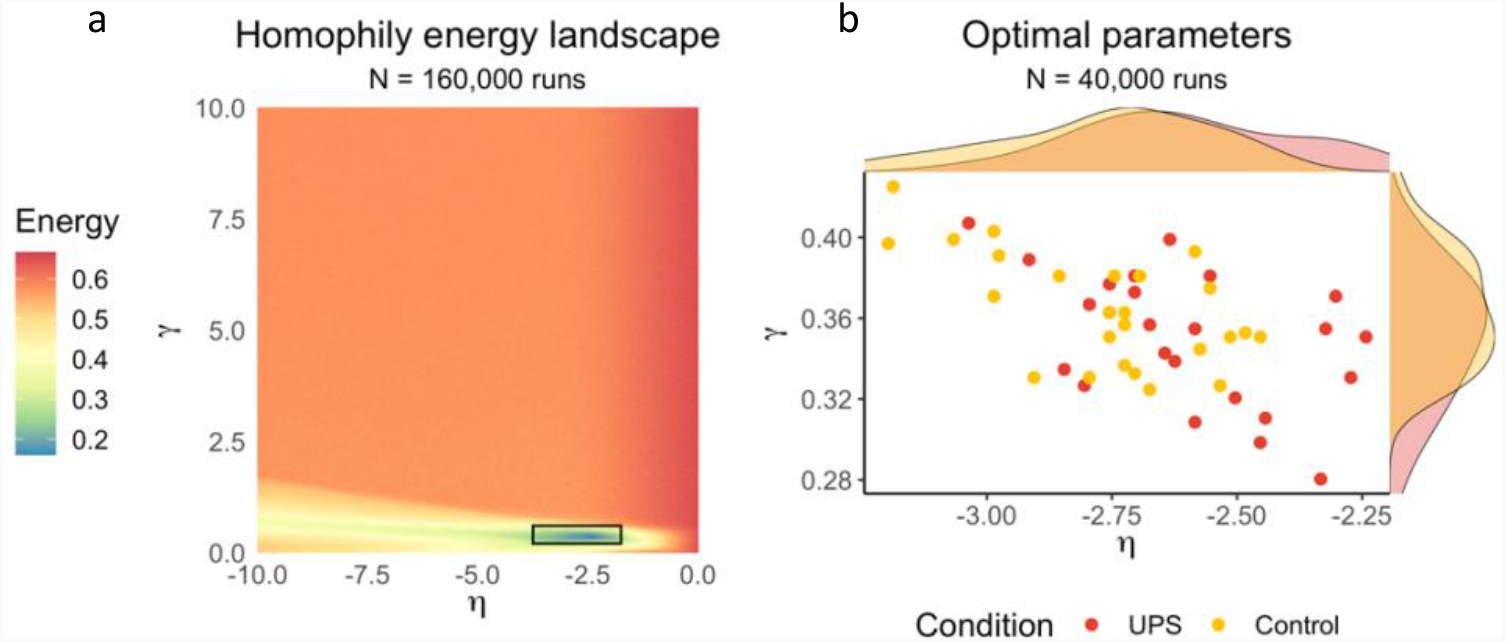
Adversity attenuates optimal generative modelling parameters. **(a)** In the first run of the homophily model, 160,000 unique combinations of cost parameter *η* and value parameter *γ* were tested. The energy surface shown is the sample average (N = 49). **(b)** Optimal values of *η* and *γ* produce the lowest-energy synthetic networks. Values were obtained by testing an additional 40,000 parameter combinations in a narrow low-energy window of the initial grid search, highlighted with a black rectangle in (a). Each data point in the scatterplot represents a single animal. Density plots above and to the right highlight differences between UPS and control conditions. Optimal parameters tend to fall closer to the origin for animals in the UPS condition (ANOVA F_1,47_ = 5.700, p = 0.021).

The parameters producing the lowest-energy networks for each animal are shown in **Figure 4b**. The cost and value parameters were moderately correlated (*r* = -0.574, *p* = 1.65 × 10^−5^), placing the simulations on an axis from the origin of the parameter space (*η* = 0, *γ* = 0). This indicates that simulations with a more severe penalty on long-distance connections usually had stronger preference for connections between regions with shared neighbours.

Along this axis, animals in the UPS condition tended to fall closer to the origin; we confirmed this observation by comparing the length of a vector from the origin to each point between groups (UPS *M* = 2.63, *SD* = 0.213, Control *M* 2.79, *SD* = 0.210; ANOVA *F*_*1,47*_ = 5.700, *p* = 0.021). The simulations for animals in the UPS condition were therefore subject to weaker constraints on the formation of connections. One possible confound here is that the models may simply perform better for one group than the other, but this was not the case: no difference in model energy was observed (UPS *M* = 0.101, *SD* = 0.015, Control *M* = 0.105, *SD* = 0.010; ANOVA *F*_1,47_ = 0.719, *p* = 0.401).

But what is the nature of this group difference in parameters? One possibility is that either or both parameters drive the change in a relatively independent manner. Alternatively, it could reflect a single underlying shift in wiring constraints that incorporates both parameters. We distinguished these alternatives using a partial least squares discriminant analysis (see **Methods**). This formally tests for the presence of underlying factors that explain the group difference in parameter combinations. There was a significant correlation between the group affiliation and the first latent variable (*r* = 0.36, *p*_permuted_ = 0.011) but not the second latent variable (*p*_permuted_ = 0.898). Both parameters of the generative model (*η* coefficient = -1.5549, 95% CI = [-1.8929 -1.2969]; *γ* coefficient = 0.1229, 95% CI = [0.0684 and 0.1858]) loaded significantly onto this component. There was no between-group difference in scores on the component (KS *D*_1,47_ = 0.308, *p* = 0.159). Thus, it seems that the observed group difference in location in the parameter space reflects a change that incorporates both wiring parameters, rather than reflecting one or two independent effects.

### Shift in wiring economy induces greater stochasticity

Simulations closer to the origin of the parameter space have greater stochasticity or randomness in the generative process^31^. To understand why this is the case, imagine that the edges in the wiring probability matrix are competing with one another. When the cost penalty and topological preferences are strong, fewer edges have high probabilities of wiring and the preferred winner is clear. But when constraints are weaker, more edges qualify as good contenders, giving the probabilistic nature of the process a greater role in the gradual organisation of the network.

Simulations for the UPS condition showed a flatter distribution with a greater dispersion of values in the probability matrix compared to the control condition (**Figure 5a;** KS *D*_1,47_ = 0.055, *p* = 2.20 × 10^−16^), corresponding to more potential connections with higher probabilities of wiring and therefore heightened stochasticity. Variance among wiring probabilities rose over the course of the development of each simulation, particularly in the UPS condition, indicating that this increase in stochasticity was more pronounced later in the generative process (**Figure 5b**). At the end of the generative process, the simulations for mice in the UPS condition exhibited a distribution of node degree that was closer to normal (kurtosis: KS *D*_1,47_ = 0.475, *p* = 0.005), indicating that the shift in wiring probabilities subtly randomized network topology.

**Figure 5.**
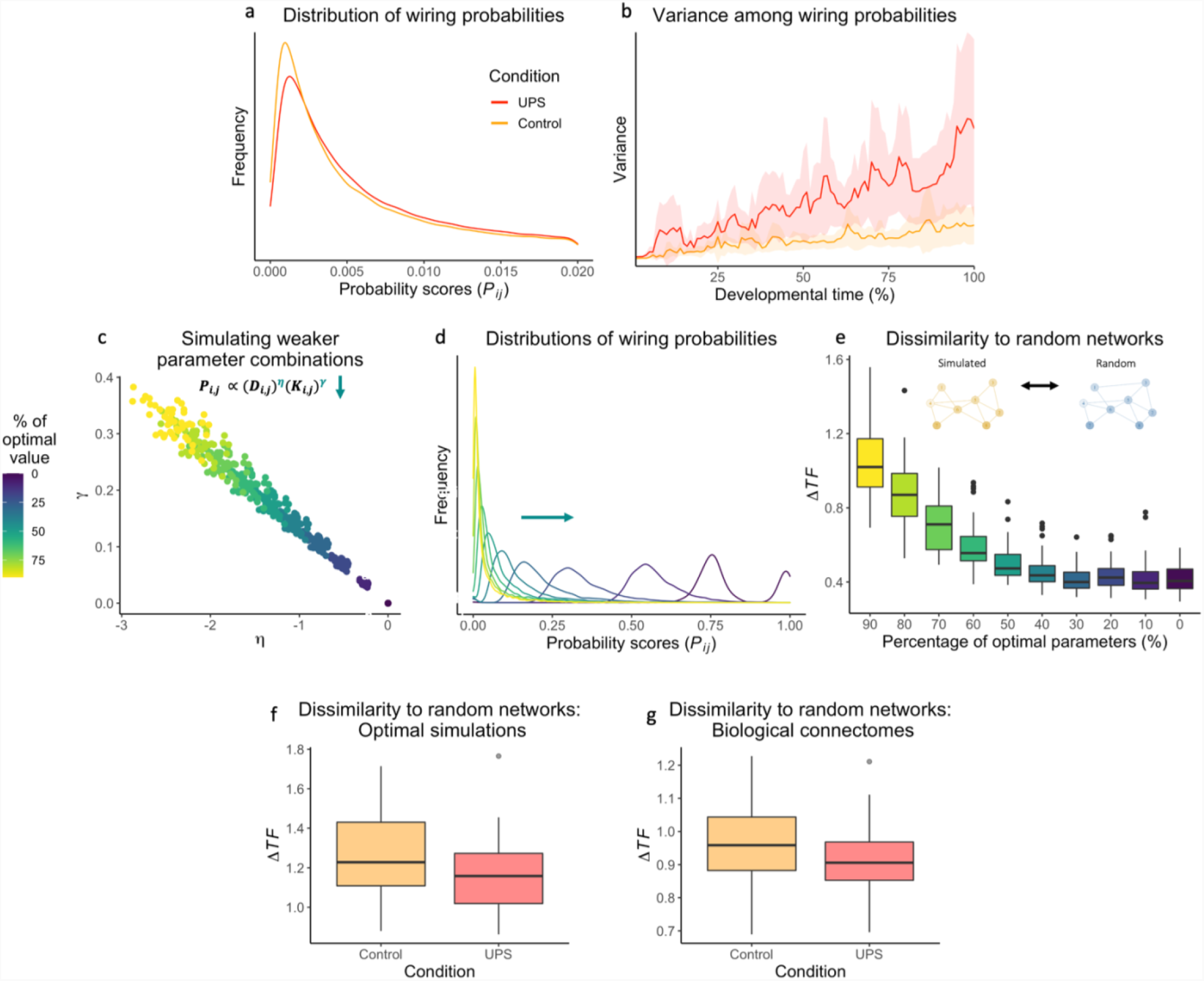
Weaker wiring constraints heighten stochasticity of network development. **(a)** Distributions of wiring probabilities (*P*_*i,j*_) within the probability matrix, taken as the group averages across all steps of optimal simulations. The UPS condition shows a flatter distribution with greater dispersion, corresponding to more connections with higher wiring probabilities. **(b)** Variance among values in the probability matrix (*P*_*i,j*_) corresponds to the dispersion of likelihoods of potential future connections. Wiring probability variance rises as simulations develop, especially in the UPS condition, indicating that model stochasticity was more pronounced later in the process. **(c)** To assess the effect of systematically manipulating wiring constraints, simulations were run at 10% increments from the optimal values for each animal to zeros. This resulted in the 490 parameter combinations plotted in this space. **(d)** Distributions of wiring probabilities (*P*_*i,j*_) within the probability matrix, taken as the average across all steps, at each parameter interval. Wiring probabilities for simulations with weaker parameters approach a normal distribution. **(e)** Topological dissimilarity (Δ*TF*; see Methods) was averaged across 1000 randomly wired networks. The organisation of simulated networks gradually resembles random topology as parameters approach zero. The same trend is observed when comparing the UPS condition to the control condition, both for **(f)** optimal generative models and **(g)** biological connectomes derived through tractography.

To explore the relationship between model stochasticity and parameters more systematically, we produced simulations using incrementally smaller values of *η* and *γ*. Specifically, we ran generative models at 10% increments from the optimal parameters for each animal, at each stage moving towards the origin of the parameter space (**Figure 5c**).

As the parameters neared *η* = 0 and *γ* = 0, the distribution of values within the wiring probability matrices (*P*_*i,j*_) exhibited greater dispersion (**Figure 5d**). This corresponds to a greater number of connections with high probability of wiring over the course of the generative process. Simulations with smaller wiring parameters had a more random topology (**Figure 5e**), as measured by the average Δ*TF* to 1000 randomly wired networks. We found the same trend toward random network topology in the UPS group, both in the optimal simulations (**Figure 5f;** UPS *M* = 1.18, *SD* = 0.214, Control *M* = 1.27, *SD* = 0.214, ANOVA *F*_1,47_ = 2.158, *p* = 0.148) and the biological connectomes (**Figure 5g;** UPS *M* = 0.915, *SD* = 0. 125, Control *M* = 0.973, *SD* = 0.126, ANOVA *F*_1,47_ = 2.647, *p* = 0.110). Though subtle, this trend is in line with the principle that weaker wiring constraints heighten stochasticity in the formation of structural connections, thereby leading to more random brain network topology.

## Discussion

We explored the effects of early adversity on the development of the structural connectome. We deployed generative network modelling in a mouse model of unpredictable postnatal stress (UPS) to test whether adversity alters the economic trade-off that governs structural brain development. The parameters that best simulated the rodent connectomes were closer to zero for the mice exposed to UPS, resulting in greater variability in wiring probabilities and therefore stochasticity in the generative process. Thus, exposure to a chaotic and unpredictable environment appears to attenuate the constraints governing brain development such that the formation of structural connections is subject to weaker control. These results point to a crucial intermediate level of explanation for the developmental impact of early adversity.

Replicating prior work in generative network modelling, models with a topological term outperformed that based purely on distance^16,21,22,34^, and models implementing the principle of homophily produced the most realistic structural connectomes^16,21,22,34^. These findings accord well with previous research on the development of the mouse brain; wiring cost alone is not sufficient to recapitulate the complex topology of its macroscopic structural networks^17^. In our study, the neighbour rule – which favours connections between regions with a greater number of shared neighbours – produced networks that possessed not only similar statistical distributions of nodal characteristics, but also their local organisation and spatial layout. Importantly, this organisation was not hard-coded into the algorithm but emerged from the trade-off between cost and value over the course of the generative process. Our study is the first to implement the two-parameter generative model in rodents and replicate the comparative success of homophily in this species. One potential explanation for its success may be that it captures macroscopic dynamics of Hebbian learning: as regions with similar neighbourhoods are likely to experience comparable patterns of stimulation, mechanisms of activity-dependent plasticity would favour their consolidation into a structural network^37,38^.

The parameters that produced the best-fitting synthetic networks differed between mice according to their exposure to early adversity. Specifically, simulations for mice in the UPS condition were subject to a more moderate penalty on long-distance connections and a weaker preference for connections between regions with shared neighbours. The negative correlation between model parameters, in line with previously findings^22,34^ (but see also^21^), indicates that individuals negotiate the wiring economy of the brain by co-varying the two constraints. However, it is still possible that a single parameter accounts for the observed group difference. Using a partial least-squares discriminant analysis, we confirmed that a single latent factor that incorporates both the cost penalty and value term best explains the relationship between model parameters and group affiliation. As evolutionary pressures have favoured heightened phenotypic plasticity in harsh, unpredictable environments, even when this is energetically costly^39,40^, the brain may respond to early unpredictable stress by attenuating overall constraints on the formation of new structural connections. This finding is particularly important because existing measures of global organisation of brain structure do not have the granularity to detect the effects of early adversity. In other words, it appears that a generative modelling approach can capture complex and subtle outcomes of adversity by reducing many measures of neural organisation to a single latent factor, namely, the wiring economy of the brain.

As lower-magnitude wiring parameters correspond to heightened model stochasticity, early adversity appears to favour more random formation of structural connections. Given that UPS mice show impaired fear learning^28^ and weaker wiring constraints are associated with poorer cognitive abilities in children^22^, our results might therefore offer a mechanistic account for the previous finding that growing up in an unpredictable environment can hamper cognitive development^41^. However, greater stochasticity in network development may also reflect an advantageous process of adaptation, as individuals exposed to early adversity tend to show skills and abilities that are conducive to successfully navigating stressful contexts^39^. Across scales, the probabilistic development of neural tissue harnesses stochastic and noisy processes to build circuits that are robust to perturbation^42^. In an adverse or unpredictable environment, heightened stochasticity in the development of the structural connectome could be adaptive if it enables the nervous system to respond more effectively to future challenges in hostile environments^40^. This proposal is consistent with a recent finding that the connectomes of children with cognitive difficulties are more robust to random attacks on networks hubs^10^.

It is important to note that, while we have verified that the organisation of the synthetic networks replicates that of the empirical connectomes, they remain simulations. As such, our generative models do not provide conclusive evidence of longitudinal differences in neural development^18^. Future work could increase the biological complexity of the simulations in a few key ways. First, as the binarization of the connectomes is a gross simplification, a generative modelling strategy that produces weighted networks would be a welcome next step. Second, as structural neurodevelopment entails not just the formation of connections but their pruning, consolidation, and myelination^43^, models may benefit from varying rules and parameters across space and time. Additionally, models could incorporate other facts known to shape the emergence of connectivity, such as the functional identity or morphology of regions^44,45^. As UPS can have sex-specific effects on brain structure^28^, future work should test for sex differences in the wiring economy of the brain. Finally, comparing the effects of UPS to a simpler paradigm that consists only of limited bedding could reveal whether unpredictability or impoverishment is responsible for the observed shift in wiring constraints.

In conclusion, we found that unpredictable postnatal stress changes the economic conditions that govern the formation of macroscopic structural connections in the brain. Our results offer a promising and mathematically specified path toward understanding how early life adversity contributes to diversity in structural brain network organisation.

## Methods

### Animals

Thirty female BALB/cByj mice were housed in breeding cages with standard bedding, and subsequently transferred to maternity cages once visibly pregnant. On postnatal day zero (P0), litters were culled to five to eight pups and randomly assigned to dams to mitigate the effects of genetics and litter size. Of 49 total pups, 25 (13 male and 12 female) were assigned to a control group, whilst 24 (12 male and 12 female) were assigned to an unpredictable early-life stress (UPS) condition. Mice in the control group were raised with standard bedding and nesting material. Mice in the UPS group received 25% of the standard amount of bedding material, no nesting material, and were separated from their dam for one hour on P14, P16, P17, P21, P22, and P25. Additional details about the paradigm are available elsewhere^30^. After weaning on P26, all mice were group housed with standard bedding and no nesting material. All experiments received the approval of the Institutional Animal Care and Use Committee (IACUC) at Yale University and were conducted in accordance with the NIH Guide for the Care and the Use of Laboratory Animals.

### Tissue and imaging acquisition

Tissue was collected from the mice in adulthood (> P70) after the conclusion of behavioural testing unrelated to this analysis. Mice were anesthetized with chloral hydrate (100 mg/kg) and, once unresponsive, transcardially perfused using cold PBS/heparin (50 units/ml) solution followed by 10% formalin (polyScience). The mice were decapitated, and intact skulls were immersed in 10% formalin at 4°C for 24 hours, transferred to sterile 1 X PBS (pH 7.4), and kept at 4°C until imaging acquisition.

Magnetic resonance images were acquired at imaging facility of New York University using a 7-Tesla scanner equipped with a cryogenic probe for enhanced signal-to-noise^46^. A modified 3D-GRASE sequence was used with an echo time (TE) of 33 ms, repetition time (TR) of 400 ms, 100 µm isotropic resolution, two non-diffusion-weighted (b0) images and 60 images acquired at unique gradient directions with b= 5000/mm^2 47^. Additional acquisition details are available in a protocol paper^48^.

Images were corrected for noise and Gibbs ringing artefacts using MRtrix3^49–51^, displacement and eddy currents using FSL^52^, and field bias using the N4 algorithm provided in Advanced Normalization Tools (ANTs)^53^.

### Connectome construction and comparisons

For each subject, a map of brain connectivity was reconstructed using probabilistic tractography. First, unsupervised estimation of tissue-specific response functions was conducted using the Dhollander algorithm^54^. The fibre orientation distribution was then estimated using multi-shell multi-tissue constrained spherical deconvolution (MSMT CSD)^55^. Probabilistic streamline fibre tracking with second-order integration (iFOD2)^56^ was performed with whole-brain seeding until ten million streamlines were reached. Fibre tracking parameters were optimized for ex-vivo rodent tissue (step size 50 µm, curvature threshold 45°, FA threshold 0.1, minimum fibre length 0.5 mm)^57,58^.

A structural connectome was then built from each tractogram using a parcellation previously adapted from the Allen Mouse Brain Atlas (AMBA) and Allen Developing Mouse Brain Atlas (ADMBA) by Rubinov and colleagues^17^. The bilaterally symmetric parcellation consists of 41 cortical and 24 extracortical regions per hemisphere, for a total of 130 regions. Using ANTs^59^, each subject image was first registered to the AMBA template space using affine and diffeomorphic transformations, then the inverse transformation was used to project the parcellation into subject space. The number of streamlines connecting each pair of regions were counted and transformed into connectivity matrices, which were symmetrized. Self-connections were removed. To eliminate spurious connections and highlight topological variation across subjects^60^, we applied a weight-based threshold of 6100 streamlines to achieve a sparse connectome density (*M* = 3.52%, *SD* = 0.13%). Thresholded connectomes were then binarized.

Connectomes were compared on five measures of global topology: (1) number of edges; (2) total edge length, approximated using the sum of the Euclidean distances between connected regions; (3) number of long-distance edges, defined as connections that are more than two standard deviations above the mean connection length across the sample; global efficiency, calculated as the average inverse shortest path length of the network^8^; small-worldness, defined as the ratio of clustering to shortest path length compared to its random network equivalent^61^, which we obtained by averaging across an ensemble of 500 networks that were randomized whilst preserving the degree distribution.

Wherever group differences were assessed, a Shapiro test was first applied to test the normality of the distributions; normal distributions were compared using ANOVA, while others were compared using a KS test.

### Generative network modelling procedure

Synthetic networks were produced for each subject through a generative modelling procedure^16,21^. First, a seed network was constructed by identifying edges shared by all mice and selecting the strongest N = 28 so that, in line with previous work^21,22^, the seed would comprise about 10% of the final network density. The use of a seed network ensures parsimony, which is particularly important given the similarity of the rodent connectomes; see **Supplementary Figure S1** for additional details on seed network construction. A single edge at a time was then added to the seed network according to the following probability equation^16^:

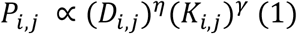

The first term *D*_*i,j*_ quantifies the distance between nodes *i* and *j*, calculated as the Euclidean distance between the centroids of each brain region in the parcellation. The parameter *η* determines the direction and strength of the contribution of distance to wiring probability (i.e., a negative value penalizes long edges while a positive value favours long edges, and a large value would produce a stronger effect than a small value).

The second term *K*_*i,j*_ quantifies the topological similarity of nodes *i* and *j* as specified by each generative rule, and the parameter *γ* determines the direction and strength of its contribution to wiring probability. As each added edge changes the topological similarity of certain nodes, *K*_*i,j*_ and *P*_*i,j*_ are continually updated at every step of the generative process. If an edge is added between nodes *i* and *j, P*_*i,j*_ is set to zero.

### Evaluation of generative models

#### Model energy

The generative process was terminated when the number of edges of the synthetic network matched that of the empirical network. The fit of each synthetic network was assessed according to the following energy equation^21^:

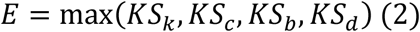

The equation consists of the Kolmogorov-Smirnov (KS) statistics comparing the synthetic and empirical networks on distributions of node degree (*k*), clustering coefficients (*c*), betweenness centrality (*b*), and edge length (*d*). Thus, a synthetic network that closely resembled the empirical connectome in all four distributions would have a low energy, while a network that greatly differed from the empirical connectome on any one of the four would have a high energy.

In addition to a purely spatial model, which did not include a topological term, we assessed three categories of generative models: two homophily models (number of common neighbours and the matching index); five clustering-based models (the average, minimum, maximum, difference in, and product of clustering coefficients); and five degree-based models (the average, minimum, maximum, difference in, and product of node degrees)^21^. Each model was computed using the Brain Connectivity Toolbox (https://sites.google.com/site/bctnet/Home) in MATLAB.

To find the optimal parameters for each model, we performed a grid search in the space defined by −10 ≤ *η* ≤ 0 and 0 ≤ *γ* ≤ 10. This approach was used to assess variability in model energies across the parameter space. A total of 160,000 parameter combinations were tested per subject and model, corresponding to 40,000 unique values of both *η* and *γ*.

#### Model topological fingerprints

To test the ability of generative models to replicate local hallmarks of empirical connectivity, we calculated the topological fingerprint matrices of both empirical and synthetic networks. TF matrices are a recently developed measure that consist of n-by-n correlation matrices of n local network statistics^31^. We included six common measures of topology in our TF matrices: node degree, betweenness centrality, clustering coefficient, edge length, local efficiency, and matching index. Each measure was calculated for all 130 nodes, then the Pearson correlation between each pair of measures was calculated and the correlations were averaged across subjects. A visual comparison of the synthetic and empirical TF matrices provides a heuristic for assessing the similarity of the correlational structure of their topology, and thereby evaluating the generative models’ ability to replicate the organisation of empirical networks. To quantify this formally, we calculated the difference in their TF matrices according to the following equation^31^:

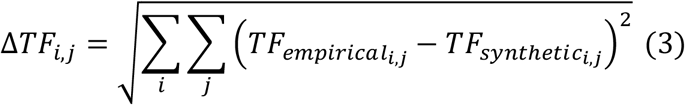

Here, Δ*TF* is calculated as the Euclidean norm of the difference between empirical and synthetic *TF* matrices. We used Δ*TF* to compare the generative rules from each category (i.e., spatial, homophily, clustering and degree) that produced the lowest-energy networks. The generative rule that obtained the lowest Δ*TF* was used in all subsequent analyses. To obtain accurate estimates of the optimal parameters for each subject, a second grid search of an additional 40,000 parameter combinations was performed in a much smaller parameter space defined by −3.75 ≤ *η* ≤ −1.75 and 0.2 ≤ *γ* ≤ 0.6.

#### Modal spatial layout

The spatial layout of the six nodal measures was also assessed^22^. Four of these measures (node degree, betweenness centrality, clustering coefficient, and edge length) are included in the energy equation while two (local efficiency and matching index) are not. For each measure, the value at each node was averaged across the synthetic networks of all 49 subjects, resulting in a single 130-by-1 vector. The same procedure was performed on the empirical connectomes. Linear correlations between synthetic and empirical vectors were then calculated. At each node, the spatial error (or discrepancy) of each measure was calculated by subtracting its average value in the synthetic networks from its average value in the empirical connectomes^22^. Thus, a lower spatial error indicates more similarity between the local topology of a particular region in the simulations and in the observed connectomes. An absolute error was calculated as the sum of the Z-scores of all six generative errors.

### Group comparisons on generative modelling parameters

A partial least squares (PLS) discriminant analysis^62^ was run to test whether the optimal model parameters (i.e. the values of *η* and *γ* producing the lowest-energy simulations) reflected of a single latent factor. The correlation between each predictor component and the primary response component was calculated, and their significance was assessed by permuting the group membership of the mice 100,000 times. For the loading of each parameter onto the PLS components, 95% confidence intervals were calculated by generating 100,000 bootstrapped samples of 49 subjects and re-computing the loadings.

The distance of each mouse from the origin of the parameter space (i.e. *η* = 0 and *γ* = 0) was then calculated and compared between groups using ANOVA.

### Exploration of model stochasticity

To explore the implications of a shift in model parameters, the composition of the wiring probability matrices (*P*_*i,j*_) was also compared between groups^31^. This was achieved by testing for differences in the distribution of probability values in the wiring matrix, taken as the average across all steps of the lowest-energy simulations. To examine whether the dispersion among probability values emerges over the course of the generative process, we calculated the variance among wiring probabilities across developmental time.

To explore the effects of attenuated model parameters more systematically, additional simulations were run scaling *η* and *γ* toward zero (i.e. running models at 90%, 80%, 70%, 60%, 50%, 40%, 30%, 20%, 10%, and 0% of the optimal parameters). The distribution of values found in the wiring probability matrices (*P*_*i,j*_) of these simulations was measured and plotted. To evaluate the randomness of simulation topology, the final networks for each of these simulations were compared to 1000 randomly wired networks using the Δ*TF* measure described above. The same comparison was conducted using the optimal simulations for each mouse, and the biological connectomes derived through tractography.

### Open access statement

Generative network modelling and analyses of synthetic networks were conducted in MATLAB, and visualisations were produced using RStudio for R. All code is available on the Open Science Framework (OSF) at osf.io/evgw5. Imaging data are available upon request to the authors. Structural connectivity matrices for each animal can be found on the OSF at osf.io/evgw5. For the purpose of open access, the author has applied a Creative Commons Attribution (CC BY) licence to any Author Accepted Manuscript version arising from this submission.

## Acknowledgements

The authors would like to thank Mikail Rubinov for providing the parcellation. This work was supported by a British Marshall Scholarship to SC, MRC intramural award G101400 to JH, MRC programme grant MC-A060-5PQ40 to DEA, and TWCF Grant 0159 to DEA. All research at the Department of Psychiatry in the University of Cambridge is supported by the NIHR Cambridge Biomedical Research Centre (BRC-1215-20014) and NIHR Applied Research Centre. The views expressed are those of the author(s) and not necessarily those of the NIHR or the Department of Health and Social Care.

## Author contributions

SC, JH, DA, and DEA conceived the analysis. AP and AK carried out the paradigm and TMA and JZ collected the imaging data. SC processed the imaging data, constructed the connectomes, executed the models, and analysed the results under the supervision of JH and DEA. SC drafted the manuscript and JH, EB, PEV, AK, DA, and DEA provided critical edits. All authors reviewed and approved the manuscript.

---

**Table S1.** Comparisons of the global topology of the empirical connectomes. See Methods for details on the computation of the connectomic measures.

| Measure | UPS<br>*M (SD)* | Control<br>*M (SD)* | Test statistic* | *p* |
| --- | --- | --- | --- | --- |
| Edge count | 296.875 (10.250) | 297.320 (11.943) | $F_{1,47} = 0.02$ | 0.89 |
| Long-distance connections | 20.583 (6.199) | 19.360 (6.945) | $D_{1,47} = 0.42$ | 0.52 |
| Maximum modularity | 0.591 (0.025) | 0.589 (0.022) | $F_{1,47} = 0.13$ | 0.72 |
| System segregation | 0.593 (0.177) | 0.496 (0.403) | $D_{1,47} = 0.26$ | 0.32 |
| Global efficiency | 0.240 (0.015) | 0.240 (0.013) | $D_{1,47} = 0.19$ | 0.71 |
| Small-worldness | 4.200 (0.395) | 4.112 (0.445) | $F_{1,47} = 0.54$ | 0.47 |

\* A Shapiro test was applied to test the normality of the distributions. Normal distributions were compared using ANOVA, while others were compared using a KS test. *Note.* “UPS” = unpredictable postnatal stress.

**Table S2.**
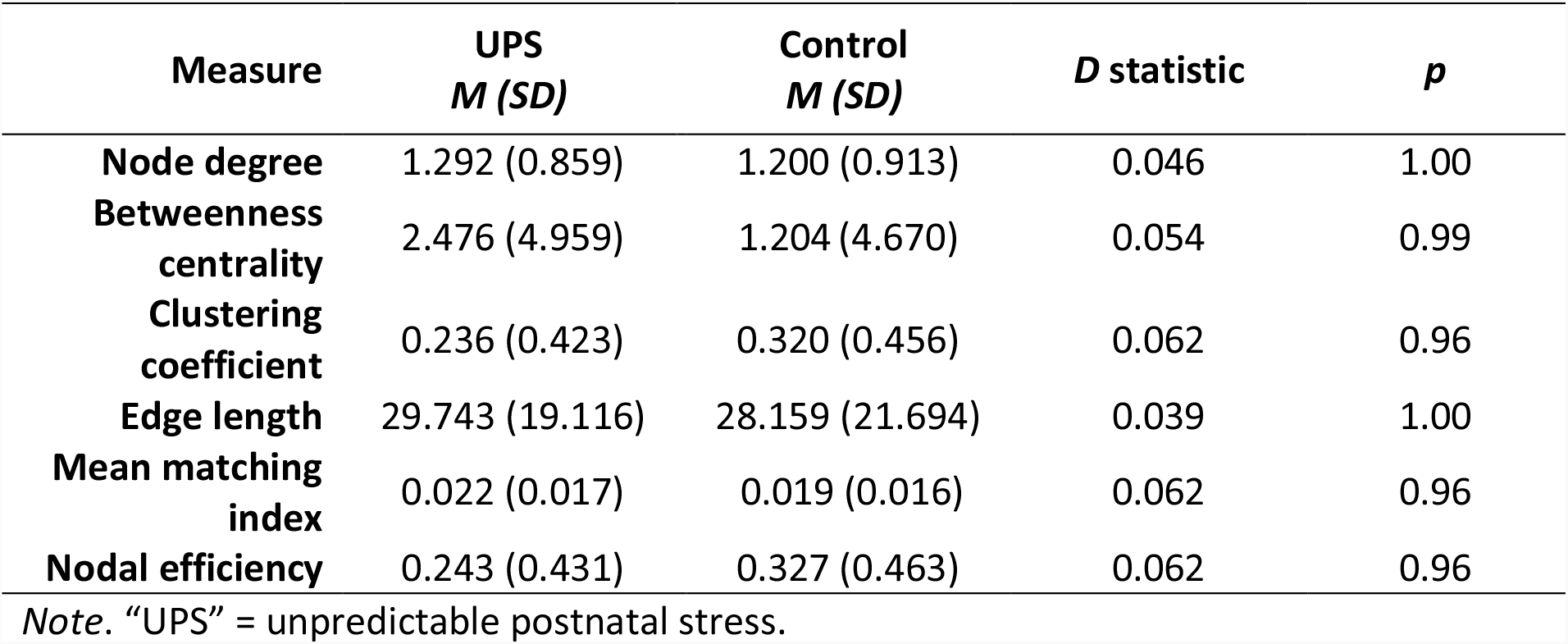
Comparisons of the local topology of the empirical connectomes. Distributions of local characteristics, taken as the group average for each node, were compared between UPS and control conditions using Kolmogorov-Smirnov (KS) tests. See Methods for details on the computation of the connectomic measures.

| Measure | UPS<br>*M (SD)* | Control<br>*M (SD)* | *D* statistic | *p* |
| --- | --- | --- | --- | --- |
| **Node degree** | 1.292 (0.859) | 1.200 (0.913) | 0.046 | 1.00 |
| **Betweenness centrality** | 2.476 (4.959) | 1.204 (4.670) | 0.054 | 0.99 |
| **Clustering coefficient** | 0.236 (0.423) | 0.320 (0.456) | 0.062 | 0.96 |
| **Edge length** | 29.743 (19.116) | 28.159 (21.694) | 0.039 | 1.00 |
| **Mean matching index** | 0.022 (0.017) | 0.019 (0.016) | 0.062 | 0.96 |
| **Nodal efficiency** | 0.243 (0.431) | 0.327 (0.463) | 0.062 | 0.96 |

Note. "UPS" = unpredictable postnatal stress.

**Figure S1.**
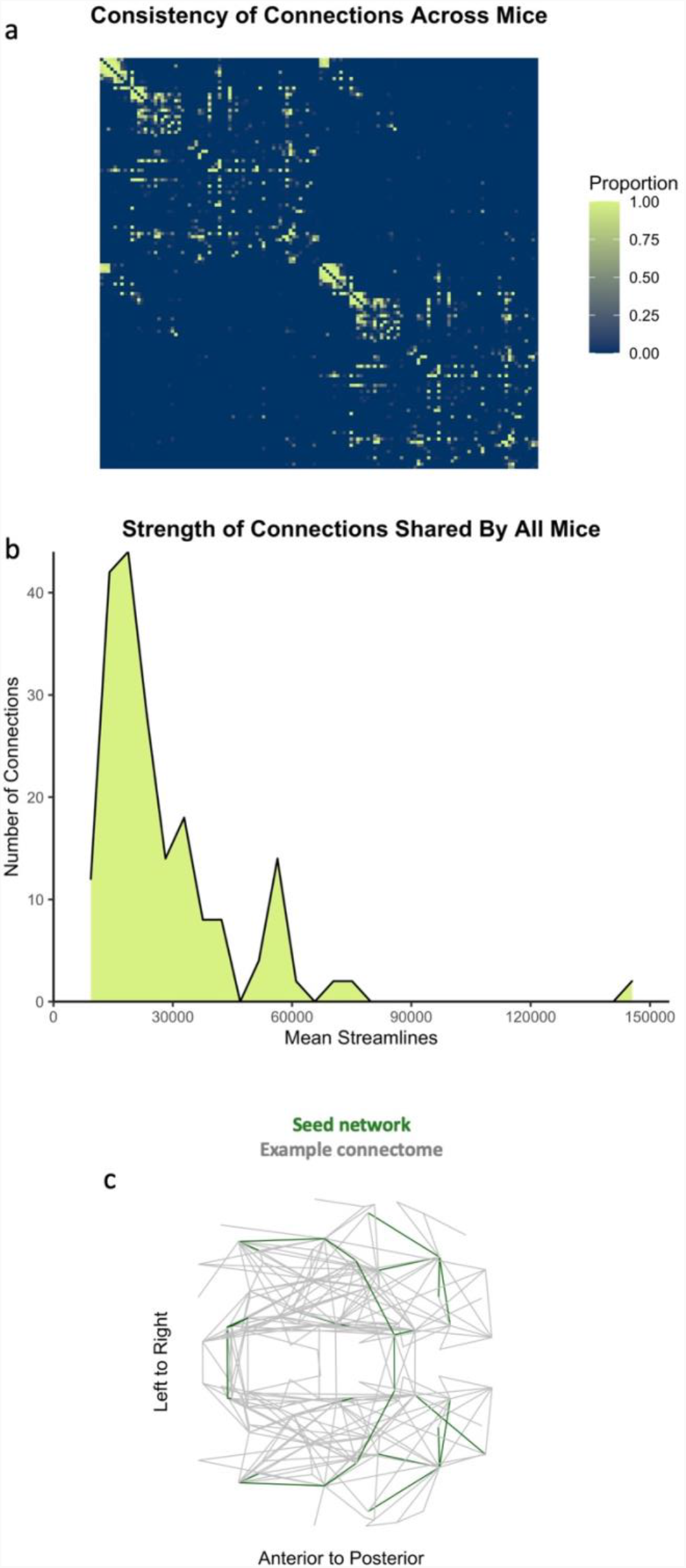
Seed network used for generative modelling. **(a)** An adjacency matrix of the 130 regions of the parcellation. The colour indicates the proportion of the sample whose connectomes contain each connection, ranging from 0 (blue) to 1 (green). **(b)** The sample mean of the weight of connections shared by all empirical connectomes. **(c)** A schematic representation of the seed network (green) superimposed over a representative empirical connectome (grey).

**Table S3.** Energy, optimal parameters, and topological dissimilarity for each generative rule. Lowest-energy networks for each animal (N = 49) were obtained by comparing 160,000 combinations of parameters in the space defined by −10 ≤ *η* ≤ 0 and 0 ≤ *γ* ≤ 10.

| Generative Model | Energy | | Eta ($\eta$) | | Gamma ($\gamma$) | |
| --- | --- | --- | --- | --- | --- | --- |
|  | Mean | CV | Mean | CV | Mean | CV |
| **Spatial** | 0.309 | 6.848 | -4.185 | -12.314 | 5.120 | 52.631 |
| **Neighbour** | 0.103 | 12.562 | -2.667 | -9.817 | 0.360 | 8.488 |
| **Match** | 0.110 | 11.276 | -2.624 | -9.842 | 0.402 | 11.236 |
| **C-Avg** | 0.170 | 11.972 | -7.301 | -13.144 | 2.361 | 34.19 |
| **C-Min** | 0.214 | 8.094 | -5.586 | -22.763 | 0.546 | 22.261 |
| **C-Max** | 0.178 | 12.184 | -8.36 | -15.669 | 4.946 | 52.494 |
| **C-Diff** | 0.207 | 10.056 | -6.166 | -8.732 | 0.880 | 24.858 |
| **C-Prod** | 0.215 | 8.176 | -5.686 | -14.933 | 0.569 | 15.848 |
| **D-Avg** | 0.098 | 9.357 | -4.787 | -13.923 | 2.596 | 12.808 |
| **D-Min** | 0.157 | 9.426 | -5.116 | -11.233 | 0.435 | 13.393 |
| **D-Max** | 0.092 | 8.806 | -4.968 | -15.855 | 2.732 | 15.693 |
| **D-Diff** | 0.132 | 10.305 | -5.514 | -25.623 | 2.507 | 21.68 |
| **D-Prod** | 0.155 | 8.889 | -5.000 | -11.119 | 0.379 | 14.452 |

*Note.* “ $\Delta TF$ ” = topological fingerprint dissimilarity, : “Neighbour” = Number of Shared Neighbours, “Match” = Matching Index, “C-Avg” = Average Clustering, “C-Min” = Minimum Clustering, “C-Max” = Maximum Clustering, “C-Diff” = Difference in Clustering, “C-Prod” = Product of Clustering, “D-Avg” = Average Degree, “D-Min” = Minimum Degree, “D-Max” = Maximum Degree, “D-Diff” = Difference in Degree, “D-Prod” = Product of Degree.

**Figure S2.**
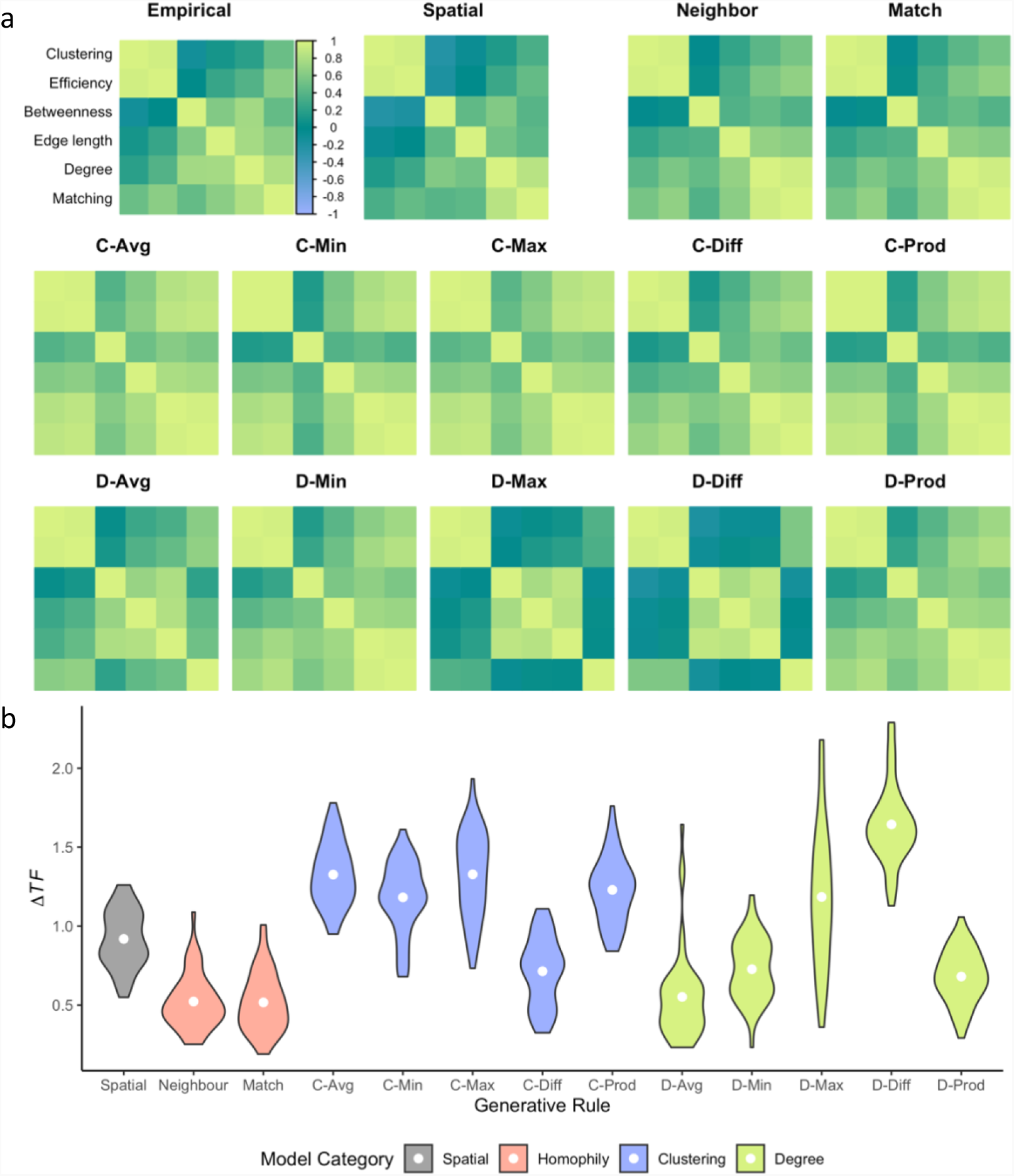
Topological fingerprint matrices and topological dissimilarity for all generative models. **(a)** The topological fingerprint is a correlation matrix of local network statistics, including node degree, clustering coefficient, betweenness centrality, total edge length, local efficiency and matching index. Across all matrices, the value of the correlation can be inferred from the colour bar, which spans -1 (lilac) to 1 (spring green). Correlations shown are the sample average (N = 49). For ease of visualisation, the measures are arranged according to the hierarchical clustering of measures in the empirical networks. **(c)** The neighbour model achieves lowest Δ*TF*, a measure of discrepancy between synthetic and empirical patterns of connectivity. White points indicate the sample mean (N = 49). *Note:* “Neighbour” = Number of Shared Neighbours, “Match” = Matching Index, “C-Avg” = Average Clustering Coefficient, “C-Min” = Minimum Clustering Coefficient, “C-Max” = Maximum Clustering Coefficient, “C-Diff” = Difference in Clustering Coefficient, “C-Prod” = Product of Clustering Coefficient, “D-Avg” = Average Degree, “D-Min” = Minimum Degree, “D-Max” = Maximum Degree, “D-Diff” = Difference in Degree, “D-Prod” = Product of Degree.

**Table S4.**
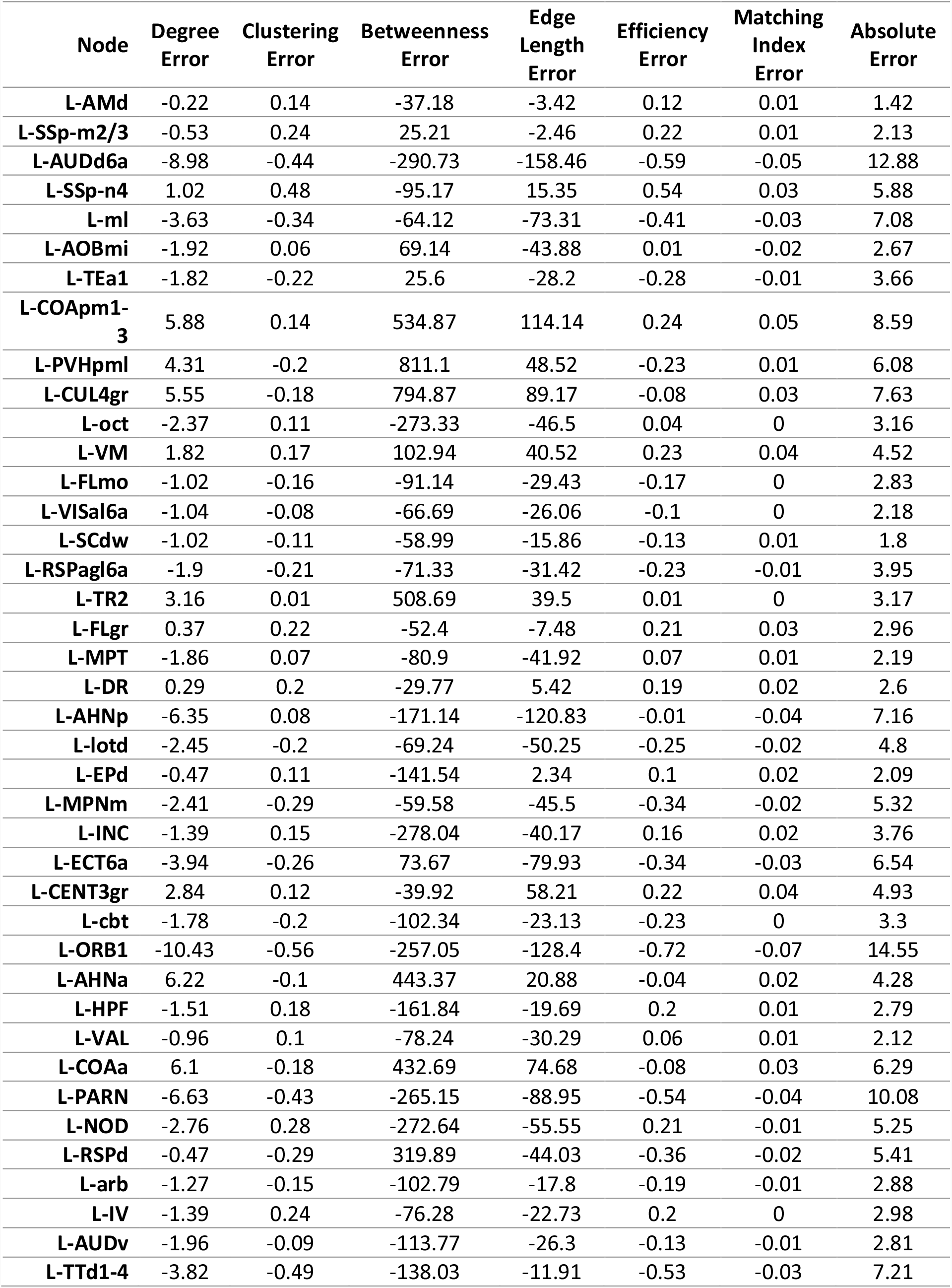
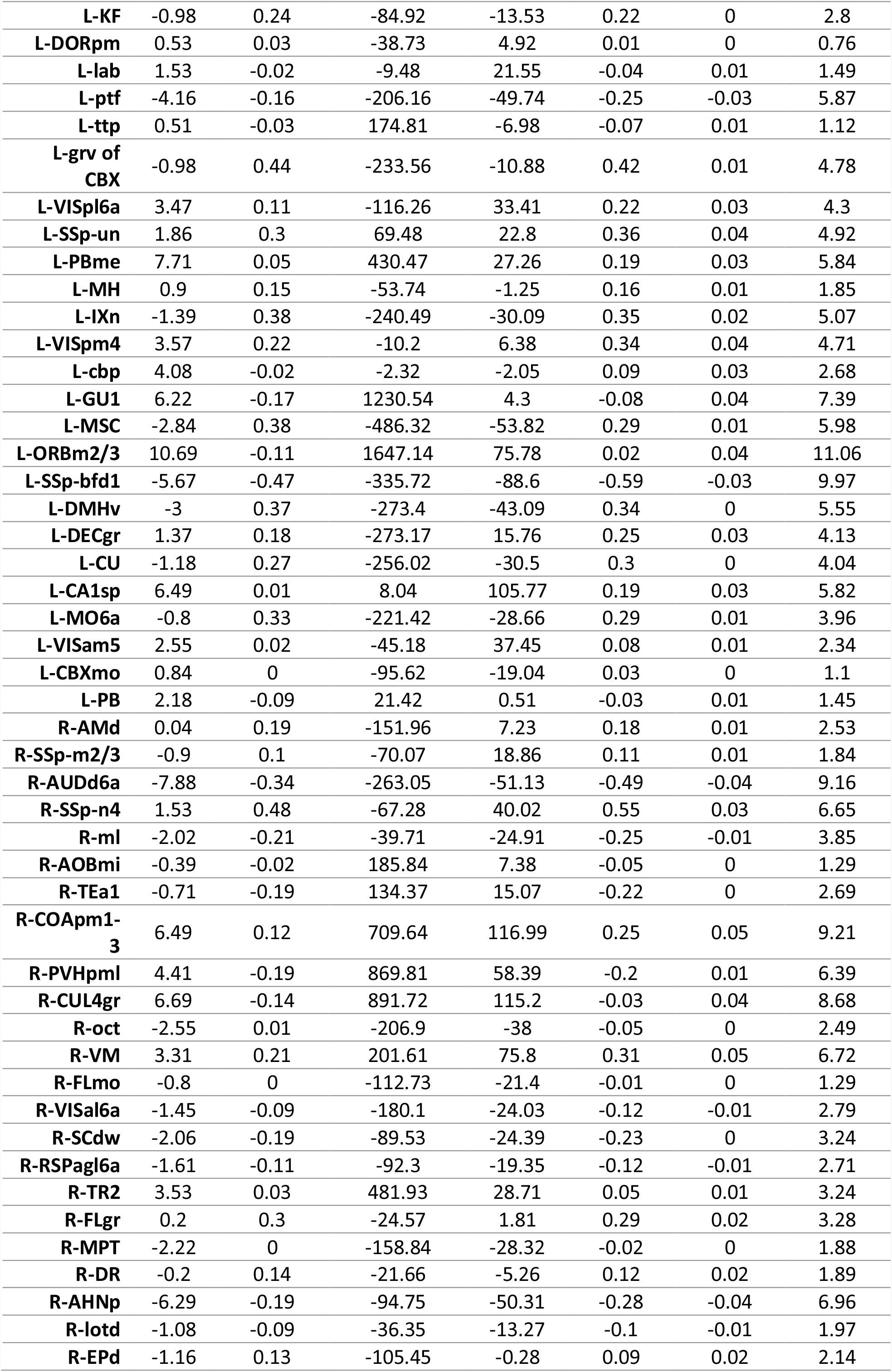
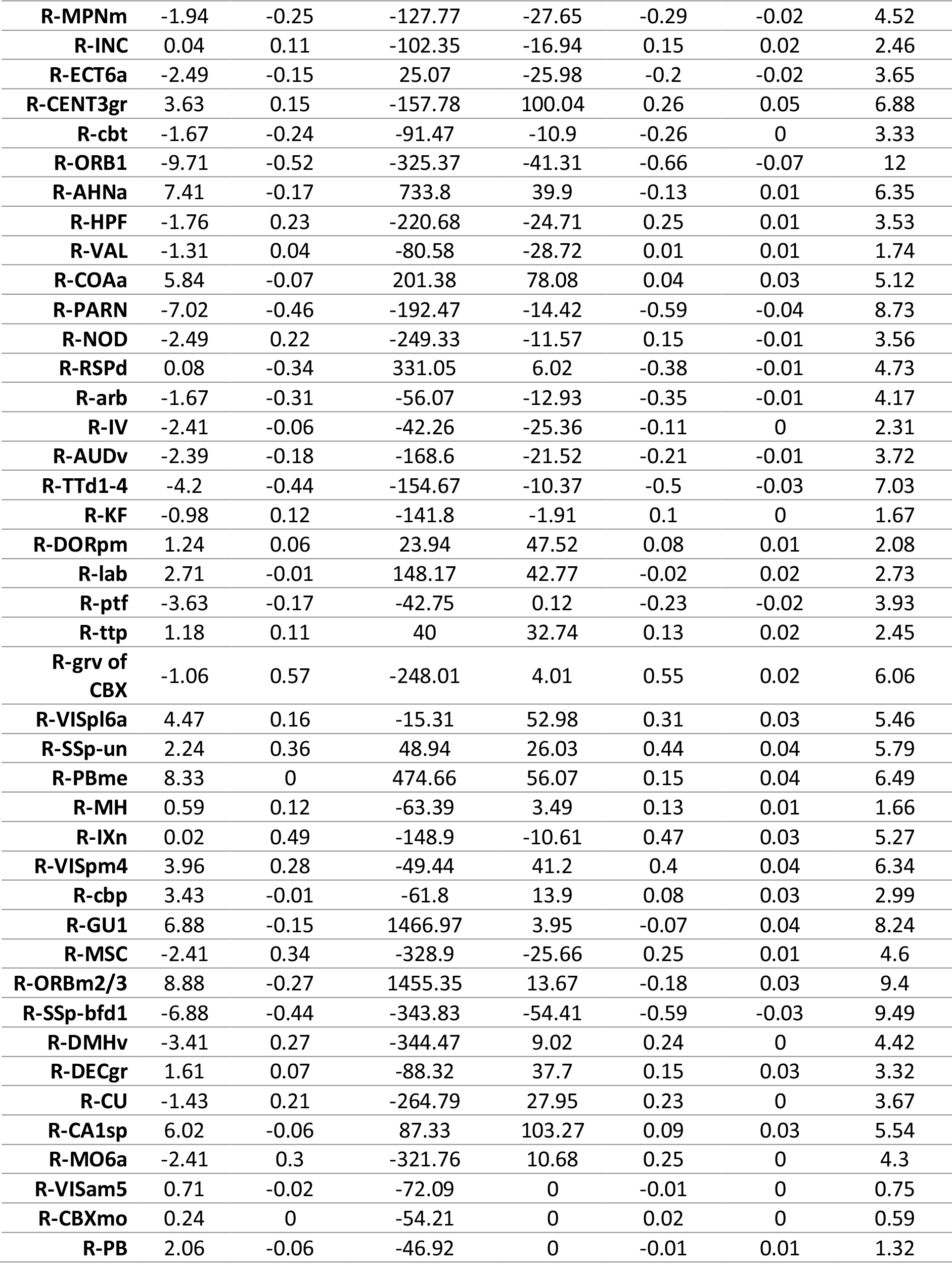
Error across topological measures and nodes. For each measure, the error quantifies the discrepancy observed between synthetic and empirical networks, while the absolute error is calculated as the sum of the Z-scores of all six errors. For details about each region, see [1].

**Table S5.** Correlations between error in spatial embedding and seed network characteristics. The error in spatial embedding quantifies the discrepancy between the synthetic and empirical connectomes on one of six nodal characteristics: node degree, clustering coefficient, betweenness centrality, edge length, nodal efficiency, and matching index. For each characteristic, Pearson correlation coefficients were calculated between its value in the seed network and the mean spatial error across the sample.

| Nodal characteristic | Correlation coefficient | *p* value |
| --- | --- | --- |
| **Degree** | 0.4356 | $2.221 \times 10^{-7*}$ |
| **Clustering** | -0.0104 | 0.9067 |
| **Betweenness** | 0.1167 | 0.1862 |
| **Edge length** | 0.1912 | 0.0293 |
| **Efficiency** | 0.0026 | 0.9767 |
| **Matching** | 0.0822 | 0.3523 |

\* Correlation is significant at Bonferroni corrected $p < 0.00833$

